# A novel multifunctional role for Hsp70 in binding post-translational modifications on client proteins

**DOI:** 10.1101/2021.08.25.457671

**Authors:** Nitika, Bo Zheng, Linhao Ruan, Jake T. Kline, Jacek Sikora, Mara Texeira Torres, Yuhao Wang, Jade E. Takakuwa, Romain Huguet, Cinzia Klemm, Verónica A. Segarra, Matthew J. Winters, Peter M. Pryciak, Peter H. Thorpe, Kazuo Tatebayashi, Rong Li, Luca Fornelli, Andrew W. Truman

## Abstract

Hsp70 interactions are critical for cellular viability and the response to stress. Previous attempts to characterize Hsp70 interactions have been limited by their transient nature and inability of current technologies to distinguish direct vs bridged interactions. We report the novel use of cross-linking mass spectrometry (XL-MS) to comprehensively characterize the budding yeast Hsp70 protein interactome. Using this approach, we have gained fundamental new insights into Hsp70 function, including definitive evidence of Hsp70 self-association as well as multi-point interaction with its client proteins. In addition to identifying a novel set of direct Hsp70 interactors which can be used to probe chaperone function in cells, we have also identified a suite of PTM-associated Hsp70 interactions. The majority of these PTMs have not been previously reported and appear to be critical in the regulation of client protein function. These data indicate that one of the mechanisms by which PTMs contribute to protein function is by facilitating interaction with chaperones. Taken together, we propose that XL-MS analysis of chaperone complexes may be used as a unique way to identify biologically-important PTMs on client proteins.

- *In vivo* confirmation of Hsp70 dimerization
- Comprehensive direct interactome of Hsp70
- Multi-domain interactions between Hsp70 and client proteins
- Identification of novel biologically-important client protein PTMs

## Introduction

The maintenance of a correctly folded proteome (proteostasis) is critical for cell survival. Cells maintain proteostasis under both basal and stress conditions through the expression of folding chaperones such as Hsp70 and its associated co-chaperone regulators (Hartl et al., 2011; Rosenzweig et al., 2019a). Hsp70 function is dependent on three conserved domains: an N-terminal nucleotide binding domain (NBD), a substrate (“client”)-binding domain (SBD), and a C-terminal (“lid”) domain (Frydman et al., 1994; Radons, 2016). The binding and hydrolysis of ATP to ADP in the NBD promotes large-scale structural Hsp70 rearrangements that allow the closing of the CTD over client proteins that bind in the SBD, promoting protein folding (Chirico et al., 1998; Rosenzweig et al., 2019b). The characterized roles of Hsp70 include folding of new and denatured proteins; transport of mitochondrial proteins and disaggregation of protein complexes (Artigues et al., 2002; Bush and Meyer, 1996; Nitika and Truman, 2017).

The essential nature of Hsp70 function in the cell, as well as its involvement in a variety of human pathologies such as cancer, has driven researchers to set out to characterize Hsp70 interactors. While great strides have been made towards this goal, these efforts have been hampered by limitations in the technologies used. For example, these past efforts have utilized affinity purification followed by mass spectrometry (AP-MS), yeast two-hybrid (Y2H) and proximity proteomics methodologies, all of which lack the ability to discriminate between direct and bridged protein interactions (Gentzel et al., 2019; Millson et al., 2005; Truman et al., 2015; Vidal et al., 1996; Zhao et al., 2005). Chemical cross-linking with mass spectrometry (XL-MS) is a powerful interactomic technique that circumvents this issue, providing information on direct interactions in protein complexes by using chemical cross-linkers (Leitner et al., 2016; Liu et al., 2015). Indeed, XL-MS studies are often complementary to the traditional structural biology methods such as X-ray crystallography, nuclear magnetic resonance, and cryo-electron microscopy (Liu et al., 2015).

Importantly, a key role for Hsp70 function is stabilization and activation of a wide range of signaling molecules including those involved in processes such as DNA damage response, cell cycle control, autophagy and nutrient sensing (Dubrez et al., 2019; Gupta et al., 2018; Truman et al., 2012; Yang et al., 2013). The Hsp70 client proteins involved in these cellular processes tend to be either highly post-translationally modified (PTMs) or regulate PTMs on other proteins. In turn, these PTMs tightly regulate a multitude of protein properties including subcellular localization, enzymatic activity and protein interactions (Nitika et al., 2020). Advances in mass spectrometry-based methods have allowed identification of more than 200 different types of PTMs on proteins including phosphorylation, acetylation, and ubiquitination (Beltrao et al., 2012; Catherman et al., 2014; Dushukyan et al., 2017). Given the numerous PTMs identified on proteins, researchers are now facing difficult choices when selecting specific PTMs for further study. Computational methods for identifying important PTMs on proteins have been partially successful but rely on pre-existing MS data (Beltrao et al., 2012; Swaney et al., 2013). In this report, we have utilized XL-MS to comprehensively understand the Hsp70 interactome. In doing so, we have uncovered not only a new set of Hsp70 client proteins, but show that these clients bind at multiple sites on Hsp70, including the N-terminal NBD. Notably, many of the Hsp70 interactions with client proteins are in close proximity to biologically-important PTMs. All in all, our data suggest that our XL-MS approach to chaperone interactome characterization can also be used as a novel way to identify biologically-important and previously unknown PTMs on client proteins.

## Results

### Analysis of cross-linked yeast Hsp70 complexes

Previous studies have identified proteins in complex with yeast Hsp70 (Ssa1) using quantitative AP-MS (Truman et al., 2015). To comprehensively identify direct Ssa1 client proteins and associated surfaces of interaction, we took a novel cross-linking proteomics approach. HIS-tagged Ssa1 was expressed in *ssa1-4*Δ, a yeast strain in which all four *SSA* (Hsp70) genes have been deleted. HIS-Ssa1 complexes, with or without cross-linking with disuccinimidyl sulfoxide (DSSO), were characterized via mass spectrometry (Figure 1A). This approach facilitated the characterization of Ssa1 complexes without competition from other native Hsp70 isoforms. Quantitative proteomics identified 1,510 interactors associated with HIS-Ssa1 in the cross-linked complexes and 1,152 in the HIS-Ssa1 complexes without DSSO-mediated cross-linking (Figure 1B). We anticipated that proteins present in the DSSO-treated complexes may consist of direct Ssa1 interactors including client proteins and co-chaperones. To distinguish direct interactors of Ssa1 from indirect interactors, we filtered our data for cross-linked peptides where at least one of the identified peptides was Ssa1. We identified a total of 363 Ssa1-containing cross-linked peptides, out of which 177 were Ssa1-client/co-chaperone crosslinks and 106 were Ssa1-Ssa1 cross-links (Figure 1C). Validating our methodology, no cross-linked peptides were observed in the control uncross-linked sample. To determine whether the cross-linking process had enriched for any particular class of protein, we performed Gene Ontology (GO) analysis of unique candidate interactors of cross-linked and control samples. This GO analysis revealed enrichment of multiple cellular functions (Figure 1D). In the cross-linked samples, proteins involved in protein folding, trafficking and cell signaling were all enriched, synergistic with the established roles of Hsp70.

**Figure 1.**
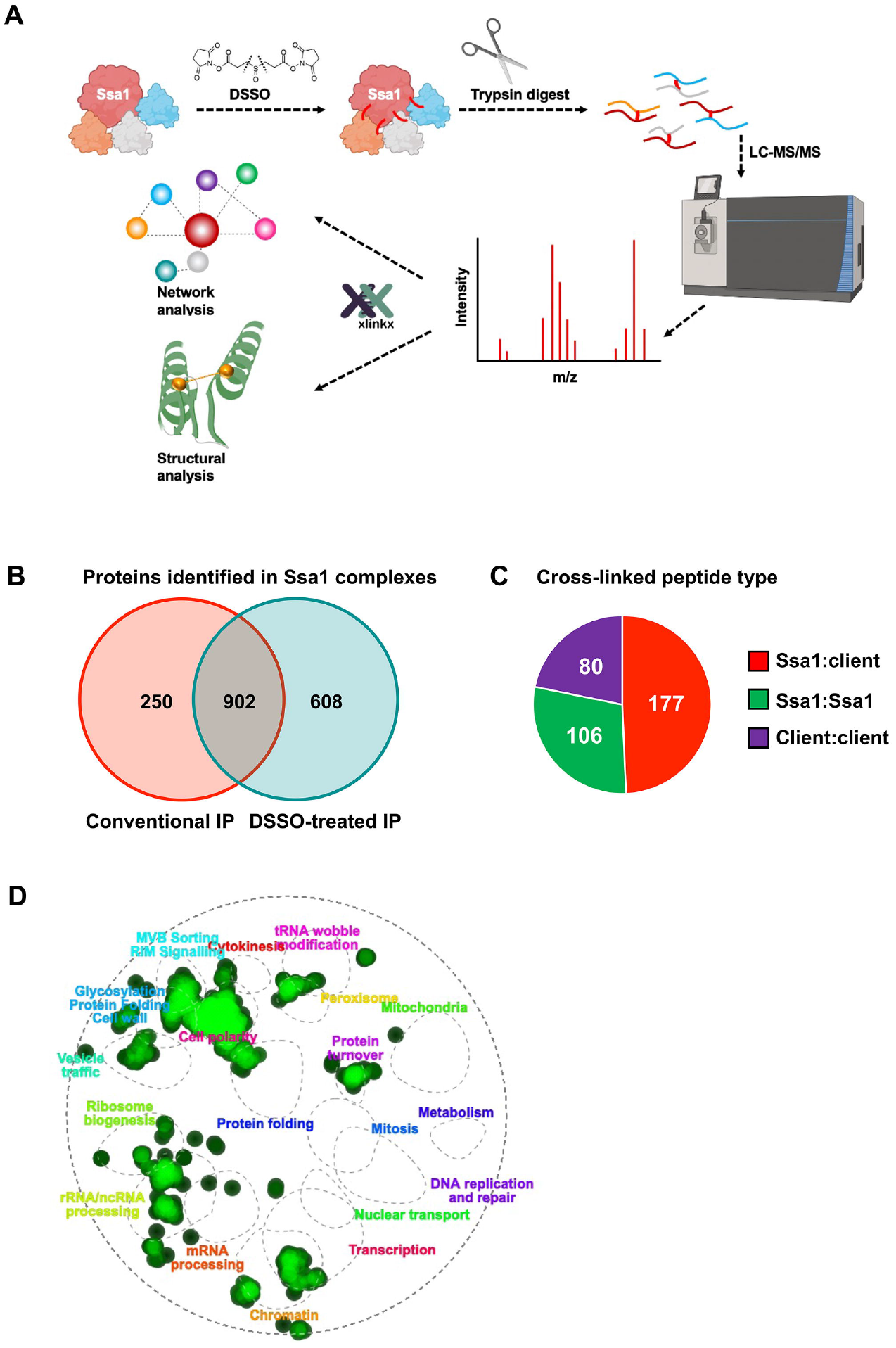
Cross-linking mass spectrometry of Ssa1 complexes. (A) Experimental workflow of cross-linking mass spectrometry of Ssa1 complexes purified from yeast cells. (B) Venn diagram representing Ssa1 complexes found in conventional IP and DSSO treated IP. (C) Pie chart showing types of cross links identified from XL-MS analysis. (D) Gene ontology analysis of DSSO treated Ssa1 immunoprecipitated complexes and crosslinked Ssa1 complexes using TheCellMap.org.

### Dimerization of Ssa1 is required for a subset of chaperone functions

Nearly a third of the cross-linked peptides detected in our experiment were between two Ssa1 peptides (Figure 1C). To distinguish whether the cross-linked Ssa1-Ssa1 peptides came from dimerized Ssa1 molecules as opposed to single intramolecular cross-links, we initially mapped the identified cross-links onto homology-based Ssa1 models. Given the large conformational change Hsp70 undergoes during its folding cycle, we utilized two models, an ADP-bound closed structure and an ATP-bound open structure model (Figure 2A and B). The mapping of cross-links onto these models revealed that a substantial number of Ssa1-Ssa1 peptides had cross-linking lengths well within the spacer arm limit for DSSO (Figure 2B), while many others exceeded the lengths possible by cross-linking within a single molecule, implying dimerization. Considering that both bacterial and mammalian Hsp70 dimerize, we evaluated whether these cross-links provided evidence for yeast Hsp70 (Ssa1) dimerization. After mapping Ssa1-Ssa1 cross-linked peptides onto a possible Ssa1 dimer structure model based on PDB structure 2KHO (Figure 2C), it was evident that at least a subpopulation of Ssa1 dimerizes in yeast (Figure 2C and 2D). To confirm this, we expressed both FLAG- and HA-tagged Ssa1 constructs in yeast. After immunoprecipitation of FLAG-Ssa1, the interaction between both tagged forms was observed (Figure 2E). Although self-interaction of yeast Hsp70 has been previously observed *in vitro* (Sarbeng et al., 2015), it has never been detected in live cells. To visualize Ssa1-Ssa1 interaction *in vivo*, we utilized bimolecular fluorescence complementation (BiFC). Yeast expressing Ssa1 tagged with Venus amino-terminal end (VN) and Venus carboxy-terminal end (VC) were examined using high-resolution fluorescence microscopy. Imaging of these cells revealed that Ssa1 dimers were clearly visible and that they localized primarily to the nucleus (Figure 2F). To demonstrate *in vivo* functionality of the Ssa1 dimer, residues identified as being present on the dimer interface based on the DnaK model (PDB: 4JNE) —E540 and N537—were mutated to A and K, respectively, generating a dimer-deficient mutant (Sarbeng et al., 2015). While yeast cells expressing the E540A/N537K dimer-deficient mutant were viable and grew at approximately WT rates, they were impaired for growth at high temperature (Figure 2G). To further explain this temperature-sensitive phenotype, we assessed the Heat Shock Response Element (HSE)-luciferase activity in WT and Ssa1 dimer-deficient cells. WT cells produced a robust HSE-luciferase signal after heat exposure and dimer-deficient cells did not (Figure 2H). Because Ssa1 is a major hub for protein folding in yeast, we set out to examine the possibility that some of the observed Ssa1-Ssa1 interactions might be the result of active Ssa1 folding a newly synthesized Ssa1 polypeptide chain. We studied the interactions of FLAG-Ssa1 (WT and substrate-binding deficient mutant V435F) with a known client, Rnr2 (Truman et al., 2015), the Ydj1 co-chaperone, and HA-Ssa1. Although WT Ssa1 co-purified with Rnr2, Ydj1 and Ssa1, the V435F mutant maintained interaction with Ydj1 and Ssa1, demonstrating that Ssa1 is not a client of other Ssa1 molecules (Figure 2I). Taken together, these findings confirm that Ssa1 dimerizes in yeast and that this self-interaction is important for a subset of Ssa1 functions.

**Figure 2.**
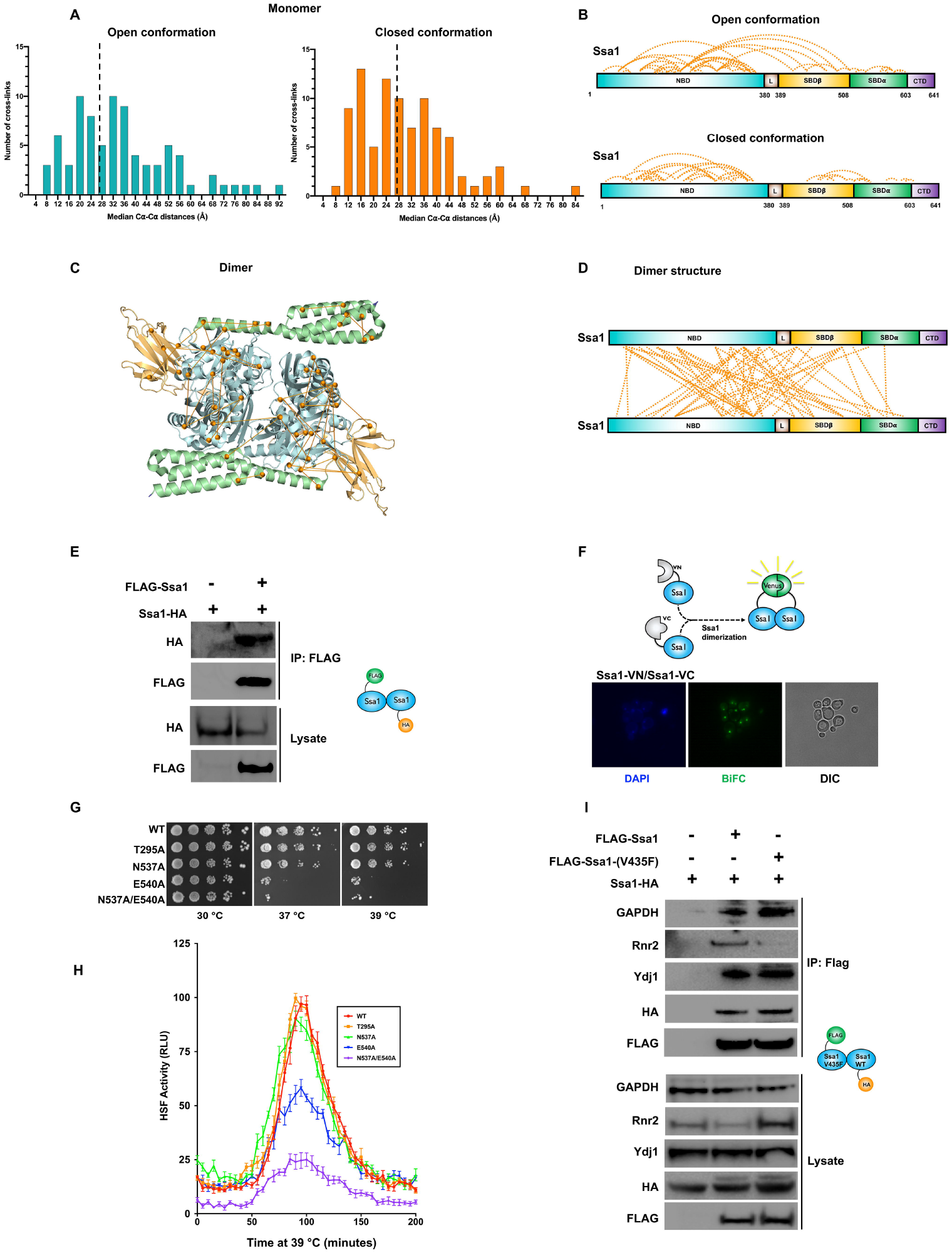
A proportion of Ssa1 exists as dimer. (A) Ssa1 cross links identified from XL-MS analysis mapped on the monomeric structure of Ssa1 in open and closed conformation. (B) Ssa1 cross links mapped on the domains of Ssa1 in open and closed conformation. (C) Internal and External Ssa1 cross links were mapped on the dimeric structure of Ssa1 (4JNE). (D) Internal and External cross links were mapped on the crystal structure of Ssa1 (4JNE). (E) Immunoblot analysis of Flag-tagged Ssa1 purified from cells expressing HA-tagged Ssa1. (F) Fluorescence images of diploid cells expressing the N-terminally VN- and VC-tagged Ssa1. DAPI was used as a nuclear marker. Scale bars are10 µM. (G) Western blot analysis of Flag-tagged Ssa1 and Flag-tagged Ssa1-V435F mutants purified from cells expressing HA-tagged Ssa1.

### Hsp70 interacts with clients throughout its domains

Our Hsp70 cross-linking strategy identified 124 new direct interactors of Ssa1 (Table S1). Unique direct binding proteins identified using XL-MS were mapped on the domain structure of Hsp70 (Figure 3A). We detected interactions on 58% of the DSSO-accessible lysines (Figure 3A). Interestingly, in contrast to the established paradigm that Hsp70 client proteins bind and interact solely at the SBD, the majority (79%) of the identified direct interactions mapped to the NBD (Figure 3A). Given that DSSO cross-links lysines, we considered that an explanation for such a high number of interactions with the Ssa1 NBD may be explained by the number and distribution of lysines present in each domain. Even when accounting for the relatively large number of cross-linkable lysines, the NBD bound over six times the number clients per cross-linkable lysine compared to the SBD (Figure S1B and S1C). While many of the NBD direct interactors relate to expected cell processes such as translation, chromatin organization and protein folding, there are several with unknown biological functions (Figure 3B). To validate our XL-MS screen, we confirmed several of our hits using co-immunoprecipitation and immunoblotting. Consistent with our MS data, Cct8, Pcl7, Ura8 and Sse1 all co-purified with Ssa1 and associated chaperones/co-chaperones Sse1, Hsp82 and Ydj1 (Figure 3C).

**Figure 3.**
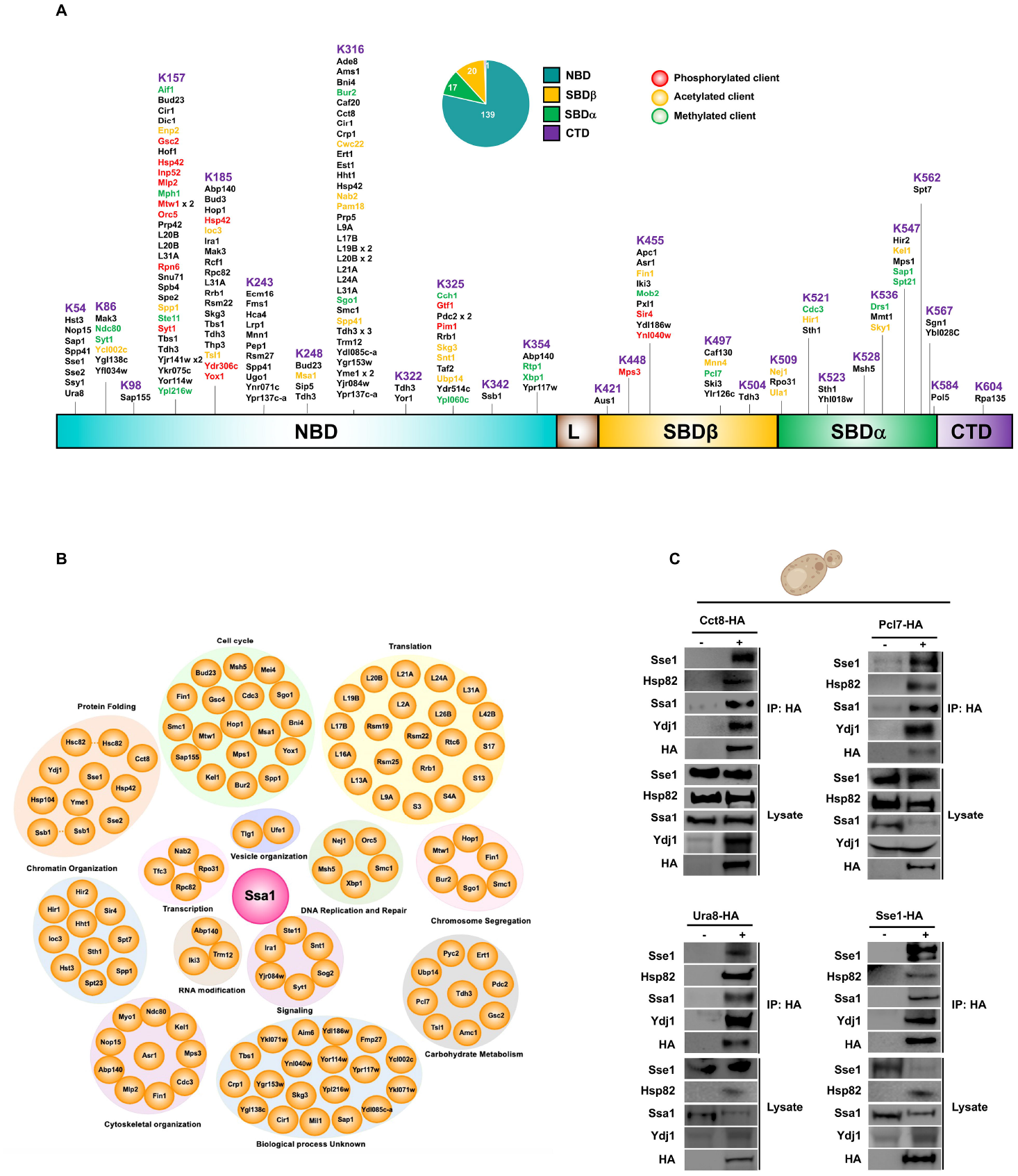
Novel clients and post translationally modified clients identified on yeast Hsp70 based on XL-MS. (A) Schematic representation of 177 inter-protein crosslinks and identified post-translational modifications on domains of Hsp70. (B) Functional classification of direct Hsp70-client peptides. (C) Western blot analysis of HA-tag immunoprecipitated Cct8, Pcl7, Ura8 and Sse1 from yeast cells.

### Exploring the biological importance of novel XL-MS-identified PTMs

31% (55/177) of our cross-linked peptides contained a PTM such as acetylation, methylation or phosphorylation (see Table S1). A search for these PTMs using GPMDB (https://gpmdb.thegpm.org/) revealed that 95% of these PTMs had not been previously observed. After considering the possibility that these PTMs might be biologically important, we selected 3 different PTM-modified cross-links for further study from proteins in diverse cellular pathways; Pim1 (mitochondrial proteostasis), Mtw1 (chromosome segregation) and Ste11 (pheromone and osmotic stress response).

### HIR complex is novel client of Hsp70

The histone regulator (HIR) protein complex regulates histone gene transcription, nucleosome formation and heterochromatic gene silencing (Mazzoni et al., 2005; Sharp et al., 2005). Our XL-MS analysis revealed a novel direct interaction between Ssa1 and HIR complex components Hir1 and Hir2. We observed cross-linking between the SBD of Ssa1 and residues K435 of Hir1 and K452 of Hir2, adjacent to their respective nuclear localization signals (Figure 4A and 4B). To validate our XL-MS finding, we carried out Co-IP and immunoblotting of Hir1 and Hir2 with key chaperone components Ssa1, Sse1, Hsp82 and Ydj1. These experiments confirmed a strong association between HIR and the chaperones tested (Figure 4C). Furthermore, we confirmed that Hir1 and Hir2 are *bona fide* Ssa1 client proteins by demonstrating that loss of Ssa1 function resulted in Hir1 and Hir2 destabilization (Figure 4D).

**Figure 4.**
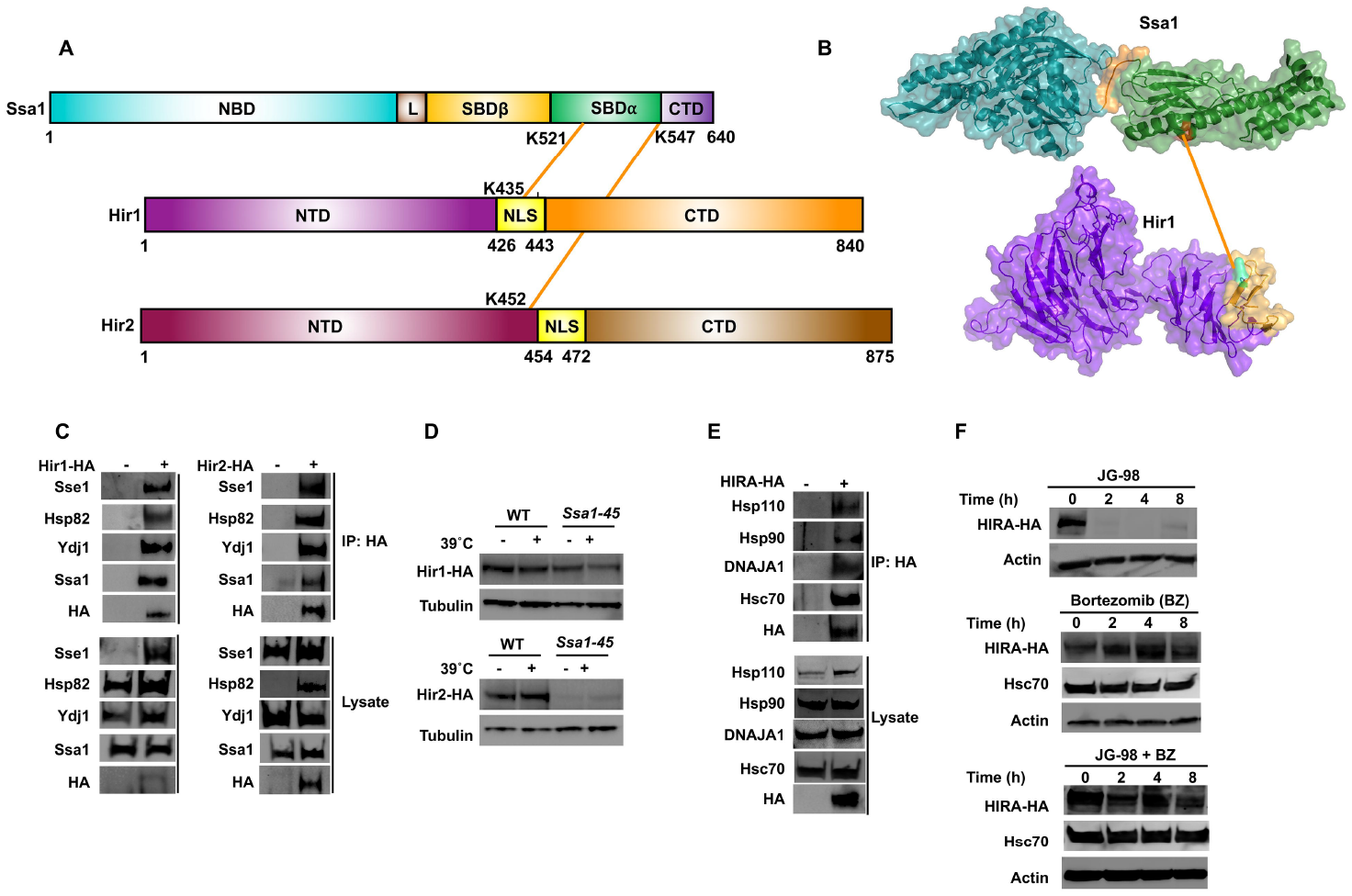
HIR complex is a novel client of Hsp70 in yeast and humans. (A) Schematic representation of Ssa1-Hir1/Hir2 inter protein cross-links detected on SBD of Ssa1 and NLS of Hir1 and NTD of Hir2. (B) Ssa1-Hir1/2 cross links mapped on the crystal structure of Ssa1, Hir1 and Hir2. (C) Hir complex interacts with the chaperone complex. (D) Hir1 and Hir2 are destabilized in Ssa1-45 mutant strain. (E) HIRA complex interacts with chaperone complexes in mammalian cells. IP analysis of the HIRA complex in mammalian cells. (F) Western blot analysis of HIRA upon addition of Hsp70 inhibitor JG-98 and proteasomal inhibitor Bortezomib.

To demonstrate evolutionary conservation of the identified chaperone-HIR interaction, we examined interaction between human Hsc70 and HIRA, the major HIR complex protein in human cells. Consistent with our results in yeast, HA-HIRA co-immunoprecipitated with Hsc70, Hsp110, Hsp90, DNAJA1 (Figure 4E). To examine dependence of HIRA on Hsc70 chaperone activity, we treated HEK293 cells with the Hsp70 inhibitor JG-98 and monitored HIRA abundance over time. HIRA levels rapidly decreased after JG-98 addition, with HIRA becoming undetectable after 2 hours (Figure 4F). Given that in our system HA-HIRA was expressed under the constitutive human cytomegalovirus (CMV) promoter, we hypothesized that the effect we observed on HIRA abundance could be explained by protein degradation. Supporting this hypothesis, addition of the proteasomal inhibitor bortezomib prevented JG-98 dependent HIRA loss (Figure 4F).

Taken together, our results suggest that HIR complex proteins are client proteins of the Hsp70 chaperone system in yeast and mammalian cells.

### Hsp70 plays a dual role in mitochondrial Pim1 protease activity

Pim1 is an ATP-dependent yeast Lon protease that is involved in degradation of misfolded mitochondrial proteins, required for mitochondrial maintenance and biogenesis (Van Dyck et al., 1994). Our XL-MS data revealed an interaction between the Pim1 protease domain and the N-terminal domain of Ssa1 (Figure 5A, 5B). We first validated the Pim1 interaction with Ssa1 and associated co-chaperones using co-immunoprecipitation and immunoblotting (Figure 5C). The clearance of mitochondrial aggregates is important for cell homeostasis and, as such, many organisms express a Pim1 homologue. To examine whether the Ssa1-Pim1 interaction is conserved in mammalian cells, we performed an equivalent experiment to that shown in 5C, using mammalian Lonp-1 as the bait. Similar to our observations in yeast, mammalian Lonp-1 interacted with chaperone proteins including Hsc70, Hsp110, Hsp90 and DNAJA1 (Figure 5D). In order to determine if Lonp-1 is a client protein of Hsp70, we treated HEK293 cells with Hsp70 inhibitor JG-98 and observed Lonp-1 degradation after 2 hours of treatment. Treatment of HEK293 cells with bortezomib before addition of JG-98 prevented loss of Lonp-1, confirming that Lonp-1 is a client protein of Hsc70 (Figure 5E).

**Figure 5.**
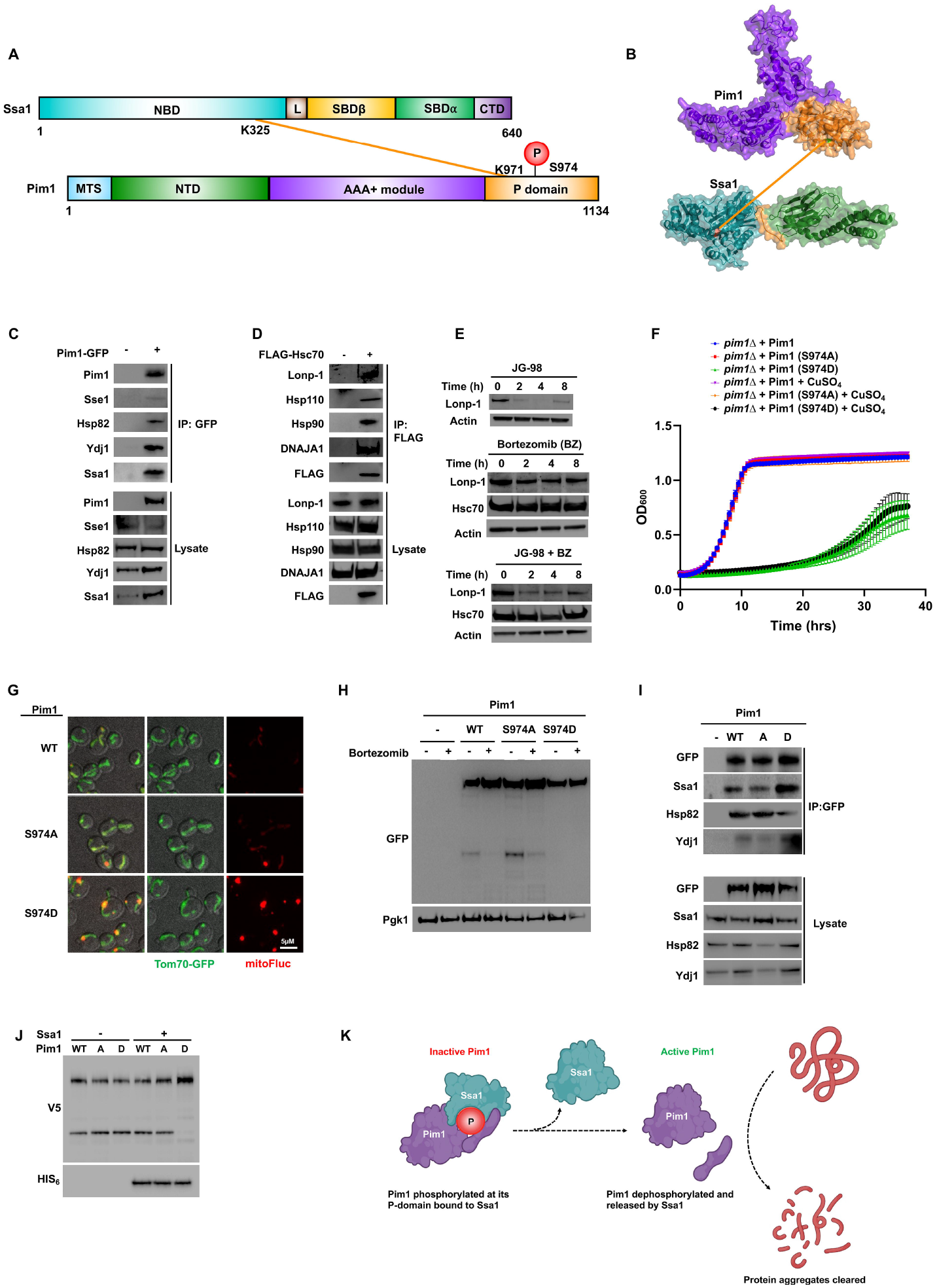
Pim1 phosphorylation regulates mitochondrial clearance in an Ssa1-dependent manner. (A) Schematic representation of Ssa1-Pim1 inter protein cross-links detected on NBD of Ssa1 and proteolytic domain of Pim1. (B) Ssa1-Pim1 cross links mapped on the crystal structure. (C) Pim1 interacts with the chaperone complex in yeast cells. (D) IP analysis of Lonp-1 interacts with chaperone complexes in mammalian cells. (E) Western blot analysis of Lonp-1 upon addition of Hsp70 inhibitor JG-98 and proteasomal inhibitor Bortezomib. (F) Growth assay of Pim1 phospho mutants in yeast. (G) Fluorescence images of cells expressing FlucSM–RFP and Tom70-GFP. Scale bars are 10 µM. (H) Western blot analysis of Pim1 wildtype and phosphomutants upon addition of Bortezomib. (I) IP analysis of Pim1 wildtype and phospho mutants with chaperone complex. (J) Western Blot analysis of Pim1 (wildtype and phospho-mutants) expressed in the presence or absence of Ssa1 *in E.coli*. (K) Schematic of Ssa1 regulation of Pim1.

The Pim1 portion of the Ssa1-Pim1 cross-linked peptide contained a previously undiscovered Pim1 phosphorylation site (S974). Given its proximity to the interaction surface between Ssa1 and Pim1, we hypothesized that this phosphorylation site might be important for Pim1 function. To examine the impact of S974 phosphorylation on Pim1 function, we expressed Pim1 mutants lacking this phosphorylation site (S974A) or mimicking constitutive phosphorylation (S974D) from the regulatable *CUP1* promoter in cells lacking Pim1. Although S974A cells grew at a similar rate to WT in standard growth media, S974D cells were substantially inhibited for growth (Figure 5F). To determine whether the growth defect of the S974D mutant was due to irregular mitochondrial protein aggregation, we examined the aggregation behavior of a previously established mitoFluc reporter (Ruan et al., 2020; Ruan et al., 2017). S974D mutants were unable to clear mitochondrial protein aggregates (Figure 5F, Figure 5G). To query whether loss of Pim1 function in the S974D mutant was due to its mislocalization, we examined localization of GFP-tagged WT, S974A and S974D Pim1 proteins. Intriguingly, Pim1 localization was unaffected by the phosphorylation state of S974 (Figure S2A).

Like many proteases, Pim1 undergoes self-cleavage to achieve full maturation and protease activity (Ondrovicova et al., 2005). We examined Pim1 processing in WT, S974A and S974D cells. In contrast to both WT and S974A, S974D resolved as a single band on SDS-PAGE, suggesting that Pim1 self-cleavage and maturation was compromised in S974D cells (Figure 5H, S2C). The proximity of S974 to the identified Ssa1-Pim1 cross-linked peptides suggested that this residue was important to the Ssa1-Pim1 interaction. Immunoprecipitation of Pim1 variants demonstrated that, although not critical for Ssa1-Pim1 interaction, S974 phosphorylation significantly enhanced the interaction between the two proteins (Figure 5I).

Based on our results shown in Figures 5A-5I, we hypothesized that Ssa1 may bind to phosphorylated Pim1 to inhibit its function. To examine Pim1 activity in the absence of endogenous Ssa1 activity, we expressed recombinant yeast Pim1 (WT, S974A and S974D) in *E. coli* (Figure 5J). In contrast to our findings in 5H, Pim1 self-cleavage was independent of S974 phosphorylation status in the absence of Ssa1 (Figure 5J). Upon co-expression of Pim1 and Ssa1 in bacteria, self-cleavage of Pim1 was restored only in the S974D mutant (Figure 5J). Taken together, our findings demonstrate that fascinatingly, Pim1 is not only a novel client of Ssa1, but that Ssa1 can prevent phosphorylation-mediated Pim1 self-cleavage (Figure 5K).

### Ssa1 regulates kinetochore function via Mtw1

Mtw1 is an essential component of the MIND kinetochore complex and it connects kinetochore subunits binding DNA to those associated with microtubules, rendering it critical to kinetochore assembly. Direct interaction between the NBD of Ssa1 and the head domain of Mtw1 was observed by XL-MS (Figure 6A, 6B). To confirm direct interaction between Mtw1 and the Hsp70 chaperone system proteins (Ssa1, Sse1, Hsp82 and Ydj1) we used a Co-IP and immunoblotting approach, similar to the one used above for other identified direct interactors (Figure 6C). Perturbation of Ssa1 function destabilized Mtw1 confirming its status as a new Hsp70 client protein in yeast (Figure 6D).

**Figure 6.**
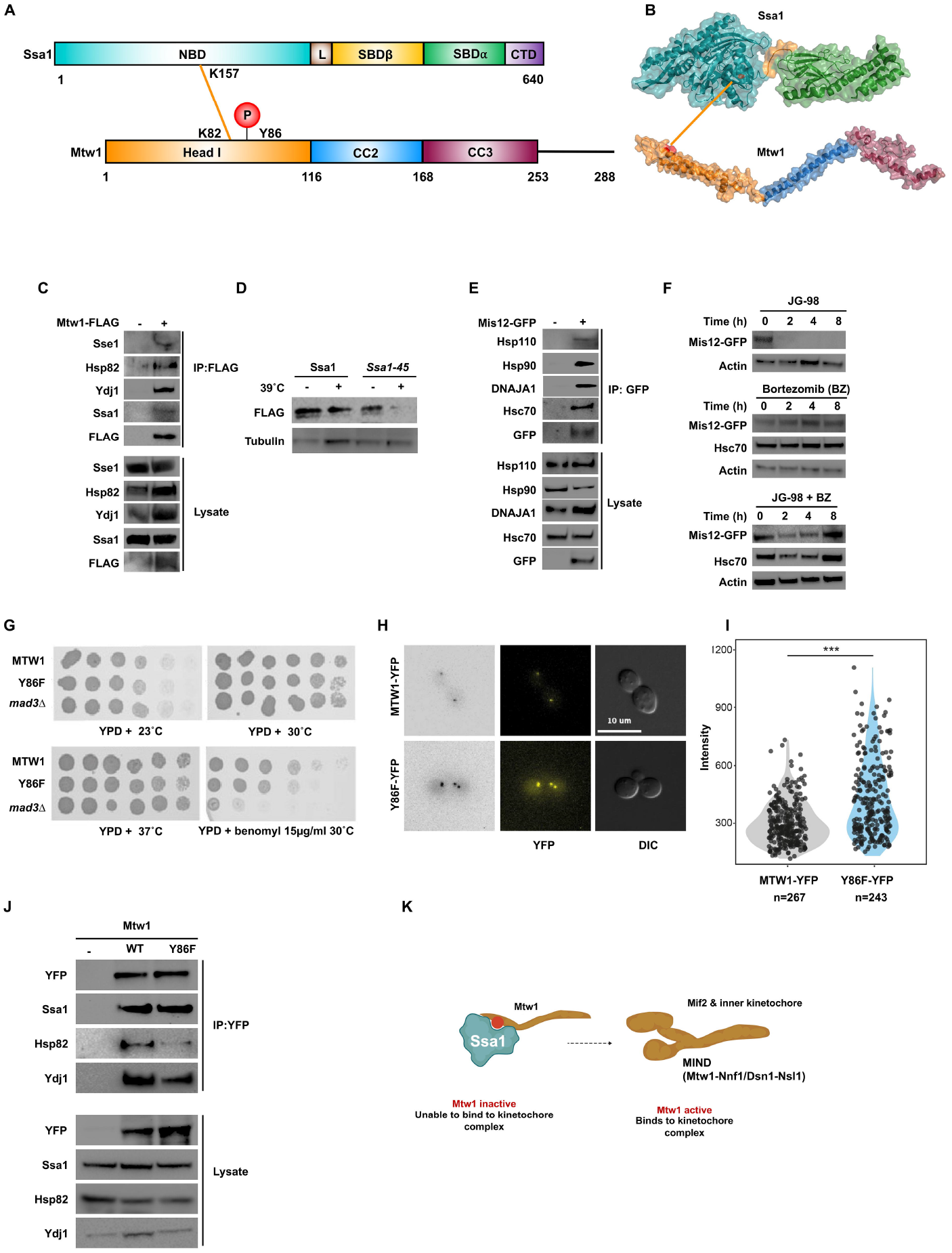
Mtw1 is a client of Hsp70 and is regulated by phosphorylation. (A) Schematic representation of Ssa1-Mtw1 inter protein cross-links detected on NBD of Ssa1 and head domain of Mtw1. (B) Ssa1-Mtw1 cross links mapped on the crystal structure of Ssa1 and Mtw1. (C) Mtw1 interacts with the chaperone complex. (D) Mtw1 is destabilized in Ssa1-45 mutant strain. (E) MIS12 interacts with chaperone complexes in mammalian cells. (F) Western blot analysis of MIS12 upon addition of Hsp70 inhibitor JG-98 and proteasomal inhibitor Bortezomib. (G) Growth assay analyzing the phenotype of the Mtw1 and its Y86 mutant. (H) Mtw1 was tagged with YFP in wild-type and Mtw1-Y86F mutant strains to compare Mtw1 localization at the kinetochore using Fluorescence microscopy. (I) Fluorescence intensities were quantified in wildtype and mutant Mtw1 using the semi-automated FociQuant ImageJ script (Ledesma-Fernandez and Thorpe, 2015). Intensities were compared using the Student’s t-test (p-value=1.8E-12). (J) Analysis of the impact of Mtw1 phosphorylation on interaction with chaperones. (K) Model of Mtw1 activity regulation via its phosphorylation.

The mammalian equivalent of Mtw1, Mis12, is critical for correct kinetochore attachment (Petrovic et al., 2010). The Ssa1-Mtw1 interaction is conserved in mammalian cells as evidenced by the successful co-purification of Mis12 with Hsc70, Hsp110, Hsp90 and DNAJA1 (Figure 6E). Furthermore, treatment of HEK293 cells with Hsp70 inhibitor JG-98 resulted in loss of Mis12, and addition of the proteasomal inhibitor bortezomib prevented JG-98-mediated Mis12 destruction, showing that inhibition of Hsp70 leads to proteasomal degradation of Mis12 (Figure 6F). Taken together, our results suggest that Mis12 is a novel client protein of Hsp70.

The identified site of interaction between Mtw1 and Ssa1 contained a previously undiscovered phosphorylation site, Y86 on Mtw1 (Figure 6A). Previous studies have shown that Y86 is at the interface between Mtw1 and a second essential kinetochore subunit, Mif2 (Dimitrova et al., 2016; Hornung et al., 2011; Killinger et al., 2020). Haploid yeast cells expressing the non-phosphorylatable Y86F Mtw1 mutant protein were viable but compromised for growth on the microtubule-perturbing agent benomyl (Figure 6G, S3A and B). Notably, we were not able to generate cells expressing a phospho-mimetic Y86E Mtw1 mutant protein. We compared Mtw1 localization at the kinetochore in both WT and Y86F mutant strains. Y86F cells showed a significantly increased accumulation of Mtw1 at the kinetochore compared to wild-type cells (Figure 6H and 6I). Although the Y86F mutation significantly impacted interaction between Mtw1 and either Hsp82 or Ydj1, Ssa1 was unaffected by the Y86F mutation (Figure 6J). Taken together our data suggests phosphorylation of Mtw1 and its interaction with Ssa1 affects its incorporation into kinetochores (Figure 6K).

### Ssa1 is involved in the selective activation of Ste11-mediated signaling

Ste11 is a MEK kinase involved in the cellular responses to both pheromone and hypo-osmolarity (Harris et al., 2001; Nishimura et al., 2016; Tatebayashi et al., 2006; Tatebayashi et al., 2020) (Figure 7A). It was an encouraging validation of our XL-MS methodology to observe direct interaction between the N-terminus of Ssa1 and the unstructured regulatory domain of Ste11, one of the first identified client proteins of Hsp90 (Figure 7B). As with Mtw1 and Pim1, the Ssa1-Ste11 peptide contained a previous undiscovered PTM, dimethylation on Ste11 R305. To determine the functional importance of dimethylation of Ste11 R305, we created the non-methylatable mutant R305A and dimethylation-mimic R305F and expressed these in cells lacking Ste11. To examine the impact of R305 on the pheromone response, we assessed the ability of Ste11 R305A and R305F to form halos in response to alpha factor, activate a FUS1-LacZ reporter and promote Fus3 phosphorylation. In all 3 experiments, both R305A and R305F behaved in a similar manner to WT (Figures 7C, D, E).

**Figure 7.**
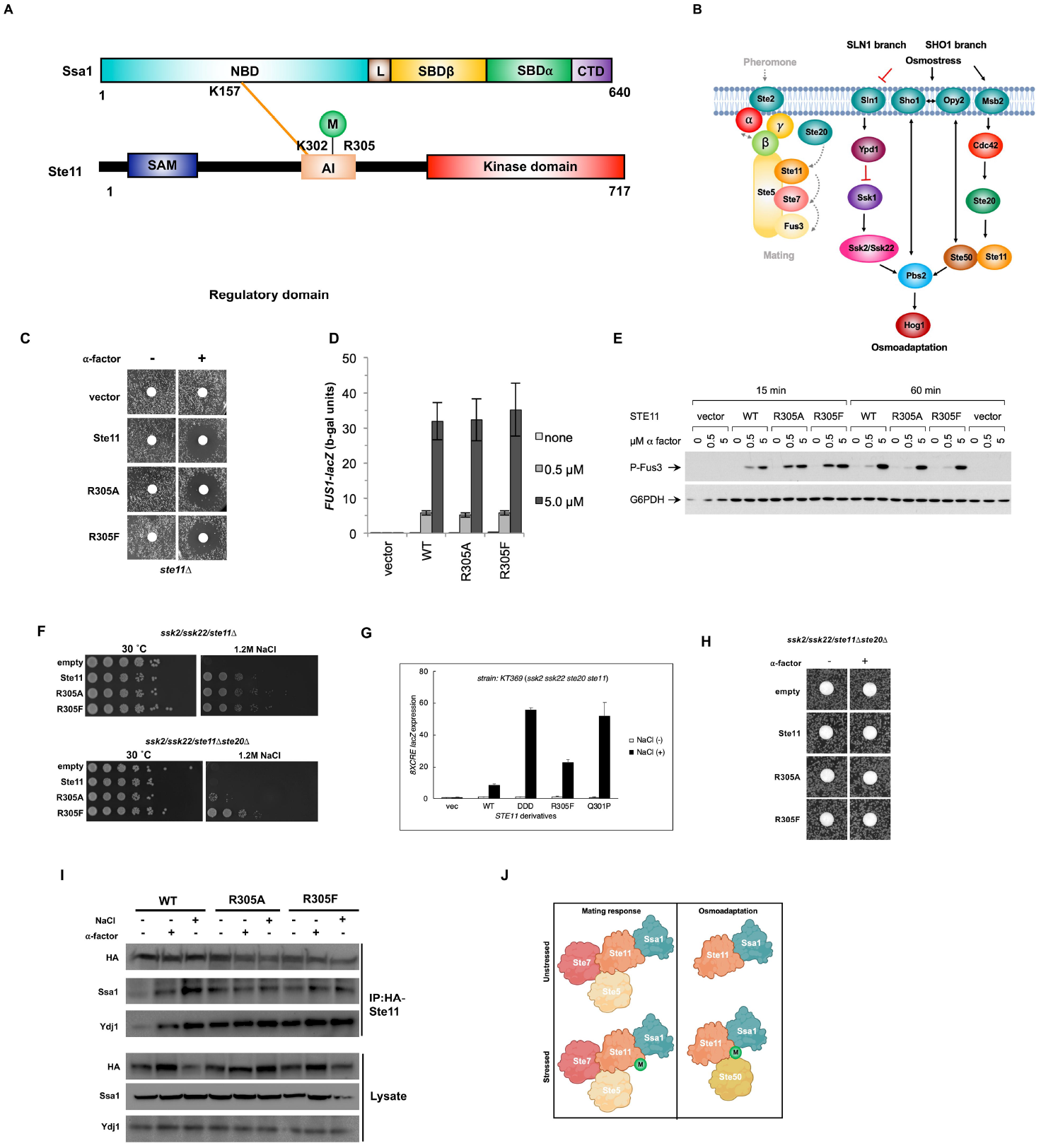
Ste11 dimethylation impacts the osmotic stress response. (A) Depiction of Ste11 pathway under hyperosmotic stress. (B) Schematic representation of Ssa1-Ste11 inter protein cross-links detected on NBD of Ssa1 and regulatory domain of Ste11. (C) Halo assay analyzing the phenotype of the Ste11 wildtype and methylation mutants in response to alpha factor. (D) FUS1-lacZ activity of Ste11 mutants in response to pheromone. (E) Western blot analysis of the effect of Ste11 wildtype and mutants in response to pheromone signaling. (F) Growth assay analyzing the phenotype of the Ste11 wildtype and the methylation mutants in hyperosmotic stress. (G) 8xCRE lacZ activity of Ste11 mutants in response to osmotic stress. (H) Halo assay showing that none of the Ste11 variants permit pheromone response in the absence of Ste20. (I) IP analysis of Ste11 with chaperone complex. (J) Model of Ste11 activity regulation via its methylation.

Although the R305 dimethylation status had minimal impact on the pheromone response, previous studies have demonstrated that some hyper-active mutations within this region (e.g. *STE11-Q301P* or *STE11-DDD*) rescue the osmo-adaptation defect caused by deletion of upstream Ste20 kinase upon high osmolarity (Tatebayashi et al., 2006; Tatebayashi et al., 2020). We examined the ability of Ste11 R305 mutants to complement the loss of Ste11 in response to osmotic shock. Because Ste11 is essential for Hog1 MAPK activation by osmostress only when the other upstream pathway (the SLN1 branch) is inactivated, we deleted the *SSK2* and *SSK22* genes encoding the MEKKs for the SLN1 branch from the host cells (Figure 7A). As with the pheromone response, the R305 dimethylation status appeared to be dispensable for Ste11 function in this regard (Figure 7F, upper panel). It is known that deletion of the upstream components of the Ste11-osmotic response pathway such as Ste20, renders cells sensitive to media containing NaCl (Nishimura et al., 2016; Tatebayashi et al., 2006; Tatebayashi et al., 2020). Interestingly, we found that while expression of WT and R305A Ste11 in cells lacking native Ste11, Ssk2, Ssk22 and Ste20 had no discernible effect on osmotic resistance, R305F Ste11 rendered cells resistant to NaCl (Figure 7F, lower panel). To determine the impact of these R305 mutations on Hog1 activation, we performed reporter assays using the Hog1 reporter *8xCRE-lacZ* on cells examined in 7F. The R305F mutant was able to induce *CRE*-mediated transcription at a level several fold greater than WT Ste11, albeit at a lower level than that observed for previously characterized hyperactive Ste11 mutants DDD and Q301P (Figure 7G). To determine whether this R305F phenotype in was specific to the osmotic stress response in ste11/ssk2/ssk22/ste20Δ cells, we performed a halo assay on the cells from 7G. In contrast to the results in 7F and 7G, Ste11 R305F was indistinguishable from WT or R305A (Figure 7H).

We wondered whether R305 dimethylation might impact interaction with either Ssa1 or Ydj1, particularly given that the Ssa1-Ste11 cross-link was formed adjacent to the R305 site. Interaction studies using immunoprecipitated Ste11 suggest that R305 dimethylation was not essential for interaction with Ssa1 or Ydj1 (Figure 7I). Taken together, our data indicates a novel interaction of Ssa1 with the unstructured regulatory domain of Ste11 adjacent to a dimethylation site that impacts only osmotic signaling (Figure 7J).

## Discussion

### Towards a comprehensive Hsp70 interactome

The identification and characterization of new chaperone interactions is important to understand the fundamental process of protein folding (Bohen et al., 1995). This knowledge can ultimately lead to the design of novel therapies that rely on the manipulation of chaperone function. While large-scale interactome studies of chaperones have been attempted previously, the methods used in these attempts have associated drawbacks that prevent a comprehensive analysis of the system (Koegl and Uetz, 2007; Yugandhar et al., 2019). For example, several of these technologies such as LUMIER are performed on purified proteins and thus might not have the dynamic range to detect the impacts of PTMs and scaffold proteins (Taipale, 2018; Taipale et al., 2014). Other cell-based assays such as AP-MS, Y2H and proximity labelling lack the ability to distinguish direct from bridged interactions. All of these drawbacks can be addressed by using the XL-MS technology we describe in this report.

XL-MS methodologies provide the ability to both stabilize transient interactions and allow characterization of the interaction surface between two proteins (Leitner et al., 2016; Liu et al., 2015). Recently, this innovative technology was used to obtain a more complete interactor list for Hsp90, providing a more accurate picture of its molecular dynamics (Chavez et al., 2016). For these reasons, we have used XL-MS to characterize the Hsp70 interactome in yeast. Our study identified a total of 1510 different proteins complexed with Hsp70, 238 of which were confirmed to be direct interactors. Importantly, 121 of these direct interactions had never been previously observed, validating the use of XL-MS technology to comprehensively study the Hsp70 interactome in the future. It is interesting to speculate what the remaining 1274 interactors represent. They may be bridged interactors present in association with Hsp70 complexes. If that is the case, it would suggest that large-scale datasets claiming to identify direct chaperone interactions might need to be revisited for accuracy. On the other hand, these interactions may be direct *bona fide* interactors that for technical reasons were unable to be cross-linked to Ssa1.

Previous *in vitro* studies suggested that the majority of interactions would be localized to the CTD of Hsp70, the domain recognized as being responsible for binding and processing of Hsp70 client proteins (Radons, 2016). Using our XL-MS method, we identified new Ssa1-interactions that map to other Hsp70 domains (Fig. 3A). Unexpectedly, 79% of the total interactions identified mapped to the NBD of Hsp70 (Fig. S1 B and C). Even after accounting for the number of cross-linkable lysines present on each Hsp70 domain, the NBD had over sixfold the number of interactions compared to the CTD. Biologically, there may be several explanations for this result. Firstly, it is possible that during the client protein-binding process, there are multiple interactions between chaperone and client-protein that engage the entirety of Hsp70. Although this phenomenon has been observed *in vitro* between recombinant Hsp70 and single client proteins, our work is the first to make a similar observation at the interactome level (Mashaghi et al., 2016). Secondly, we may be detecting the interaction of Hsp70 in fully-formed protein complexes. Finally, several of the N-terminal interactions may represent novel co-chaperones/regulators of Hsp70.

Our goal was to identify a comprehensive clientome of Hsp70. To achieve this, we did not replenish ATP during the cross-linking and purification process, skewing the complexes towards co-chaperone free, client-bound Hsp70 complexes. In agreement with this, although several co-chaperones (Sse1, Cct8, Ydj1) were detected in complexes with Hsp70, very few were identified in our cross-linked samples. Although beyond the scope of this study, future experiments may entail purification of Hsp70 complexes in different stages of the folding cycle to trap co-chaperones rather than clients.

While proteomics methods can undoubtedly produce non-native interactions, all the hits selected for follow up in this report were confirmed to be genuine Hsp70 interactors in both yeast and mammalian cells. Given the stress-dependent nature of the Hsp70 interactome, it will be important to perform variations of this XL-MS experiment under different stress conditions such as heat, cell cycle stage, DNA damage response and nutrient deprivation.

### Understanding novel Ssa1-Ssa1 interactions

An advantage of XL-MS methodologies is the ability to detect a wider range of information about protein folding and structure. For example, in the case of Hsp90, XL-MS has been used to understand protomer conformational changes upon ATP binding (Chavez et al., 2016). In this study, we identified 177 internal Ssa1-Ssa1 cross-links. Although the structure of full length Ssa1 protein has yet to be obtained, sequence similarity to bacterial and mammalian Hsp70 strongly suggests that Ssa1 forms similar ATP and ADP-bound conformations (Bertelsen et al., 2009; Rosenzweig et al., 2019b; Zhu et al., 1996). Serving as an internal control to our experiment, the majority of obtained Ssa1-Ssa1 peptides could be matched to these structures. We were surprised by the number of remaining peptides that could not be matched to any known monomeric Hsp70 structure. Deeper analysis of these peptides revealed that these were likely the result of cross-linking of *two different* Ssa1 molecules. The evidence supporting this is twofold. Firstly, the distance between these cross-linked Ssa1-Ssa1 peptides exceeded the known DSSO cross-linker length. Secondly, several of the cross-linked Ssa1-Ssa1 peptides were symmetrical; the peptides on each side of the cross-link were the same. While clearly not at the same stoichiometry as Hsp90, several studies on bacterial and human Hsp70 have demonstrated a capacity for the purified chaperone to form higher-order structures (Bertelsen et al., 2009; Liu et al., 2017; Morgner et al., 2015; Sarbeng et al., 2015; Takakuwa et al., 2019; Trcka et al., 2019) Expression of a dimerization-deficient DnaK in bacteria produces viable cells that are sensitive to thermal stress, suggesting that dimerization is needed for a subset of DnaK functions (Liu et al., 2017). Through both co-immunoprecipitation and BiFC, we demonstrate for the first time that yeast Ssa1 can also form dimers in cells. These dimers are almost exclusively localized to the nucleus, suggesting an nuclear-specific function. While the role of the Ssa1 dimer remains to be explored, given that regulation of the heat shock response by HSF occurs in the nucleus, we hypothesize that dimerization may be a novel way to regulate HSF activation in cells.

### Using sites of Hsp70 interaction to reveal novel PTMs of functional relevance on client proteins

Interactions with molecular chaperones are critical for supporting the function of proteins involved in signal transduction, particularly those involved in PTMs such as kinases or acetylases (Chen et al., 2014; O’Regan et al., 2015; Tao et al., 2016). Some of these interactions are quite stable, with client proteins requiring continuous chaperone interaction for activity (Kim et al., 2013). Others are transient, where full client protein maturation and activity requires rapid chaperone dissociation (Rosenzweig et al., 2019b). However, there is also a growing body of work that shows that PTMs play an important and novel role in chaperone interactions. For example, the client-phosphorylation status of the Hsp90-Mpk1 and Hsp90-ERK5 complexes have been shown to be important for their function (Piper et al., 2006). The chaperone code can also regulate interactions with many chaperones/co-chaperones including Hsp90, Hsp70, HSF, Cdc37 (Cloutier and Coulombe, 2013; Nitika et al., 2020; Nitika and Truman, 2017). While these PTMs have been identified on a one-at-a-time basis, ours is the first study to identify important chaperone-PTM interactions on a much larger scale.

In this work, we have followed up on three Hsp70 clients that contain novel PTMs on the site of interaction with Ssa1-Mtw1, Pim1 and Ste11. In each of these cases, while the client protein PTMs had been previously undiscovered, we have shown them to regulate novel and distinct (and very different) client protein functionalities. For Mtw1, PTM appears to regulate its localization, required for full functionality. For Pim1, S974 phosphorylation doesn’t impact stability or localization, but rather self-processing and proteolytic activity. For Ste11, one the canonical Hsp90 clients, our working model is that dimethylation regulates the transition between inactive and active Ste11 conformations, either independent of or in conjunction with its regulation by phosphorylation. While beyond the scope of this study, we hypothesize that this dimethylation is selectively impacting interactions of Ste11 with components of the osmotic stress response pathway such as Ste50.

Future studies will aim to decipher the stresses and enzymes that regulate these PTMs in addition to teasing apart their hierarchy of interaction with Ssa1. We currently have two working models; in the first the presence of the PTM recruits Hsp70 to alter client protein interactions and therefore function. In the second, Ssa1 is acting as a “protective cover” over the novel PTM, trapping the PTM in its on/off state, ultimately altering the kinetics of client activation. These two models, while reasonable, are hard to test given that Ssa1 is essential for cell growth and acts at multiple points in the signal transduction pathways involved.

This study has unveiled new regulatory mechanisms for Hsp70 and its client proteins. In doing so, we have established novel tools, methods, and workflows that will allow the field to achieve a more complete understanding of chaperones and the ways in which cells use them to integrate signal transduction pathways.

## Abbreviations

AP-MS: Affinity purification mass spectrometry
BiFC: Bimolecular fluorescence complementation
CMV: human cytomegalovirus promoter
CTD: C-terminal Domain
DSSO: disuccinimidyl sulfoxide
GO: Gene ontology
HSE: Heat Shock Response Element
HSP: Heat Shock Protein
MS: mass spectrometry
NBD: Nucleotide-binding domain
PTM: Post-translational modifications
SBD: Substrate-binding domain
VN: Venus amino-terminal end
VC: Venus carboxy-terminal end
XL-MS: Cross-linking mass spectrometry
Y2H: Yeast two-hybrid

## Acknowledgements

This work was supported by the NIH (R15GM139059 and R01GM139885 to AWT, R01GM057769 to PMP), the Queen Mary University of London and the Francis Crick Institute (Cancer Research UK—FC001183; UK Medical Research Council—FC001183 and the Wellcome Trust—FC001183 to PHT), a grant from Re-Stem Biotech to R.L., the JSPS Grants-in-Aid for Scientific Research (KAKENHI) (21H02422 to KT), the Institute for Fermentation, Osaka (G-2021-2-082 to KT). We thank Dr. Paola Lopez-Duarte for the use of her confocal microscope.

## Author Contributions

N, BZ, JET, MJW and KT conducted experiments and analyzed data with assistance from AWT and PMP. JTK, JK, RH and LF assisted with XL-MS experiments and data analysis. LR, YW and RL, contributed to the Pim1 mutation and microscopy analysis. MTT, CK and PT performed the Mtw1 microscopy. N and AWT designed the study, interpreted the data, and wrote the first draft of the manuscript. VAS edited and revised the manuscript. AWT supervised the project. All authors reviewed and approved the final manuscript.

---

**Supplementary Figure S1.**
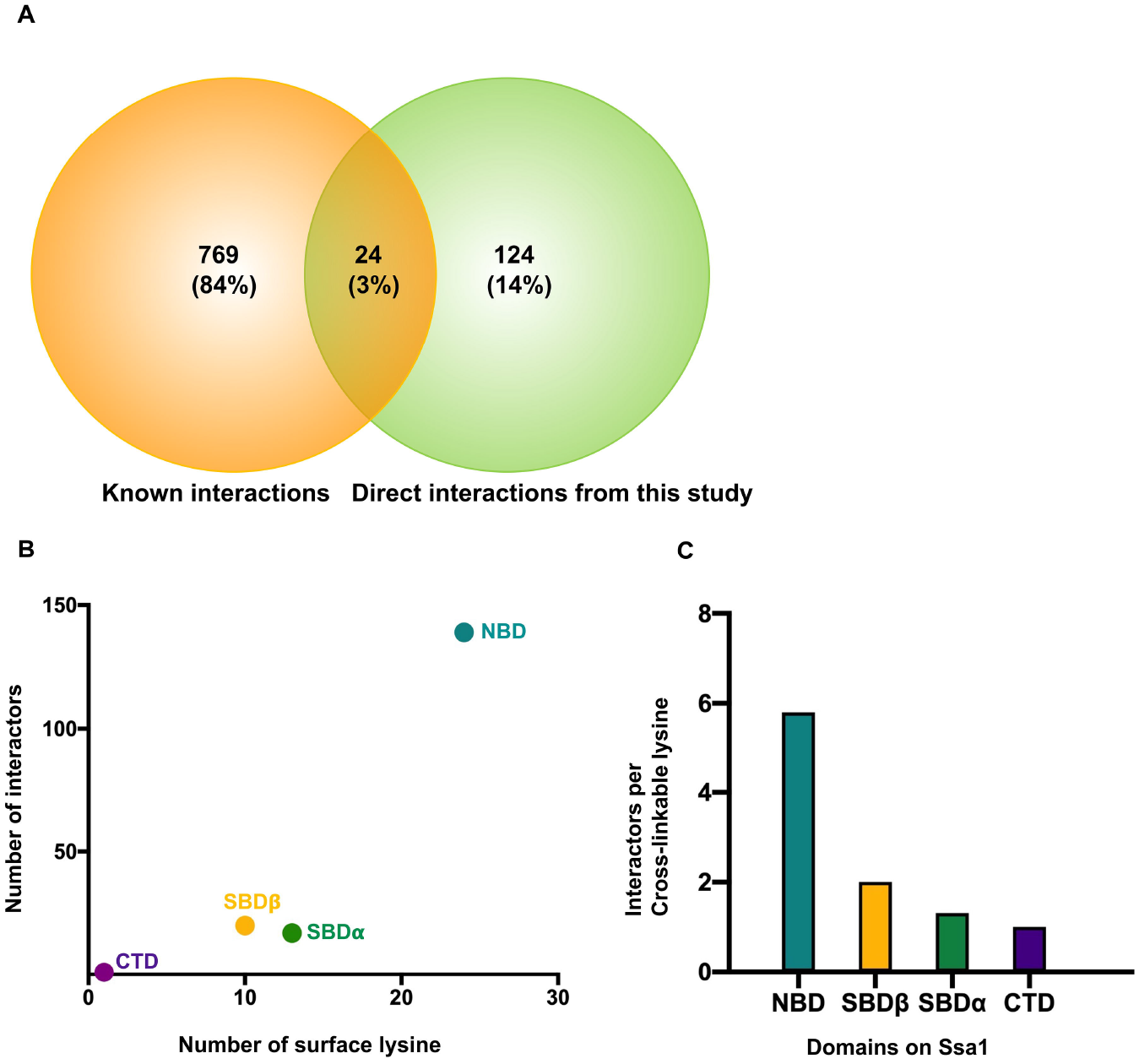
(A) Venn diagram representing previously known physical interactors of Ssa1 versus direct interactors of Ssa1 identified in this study. (B) Scatter plot of number of interactors identified versus surface lysine on the domains of Ssa1. (C) Bar graph representing interactors per cross linkable lysine on domains of Ssa1.

**Supplementary Figure S2.**
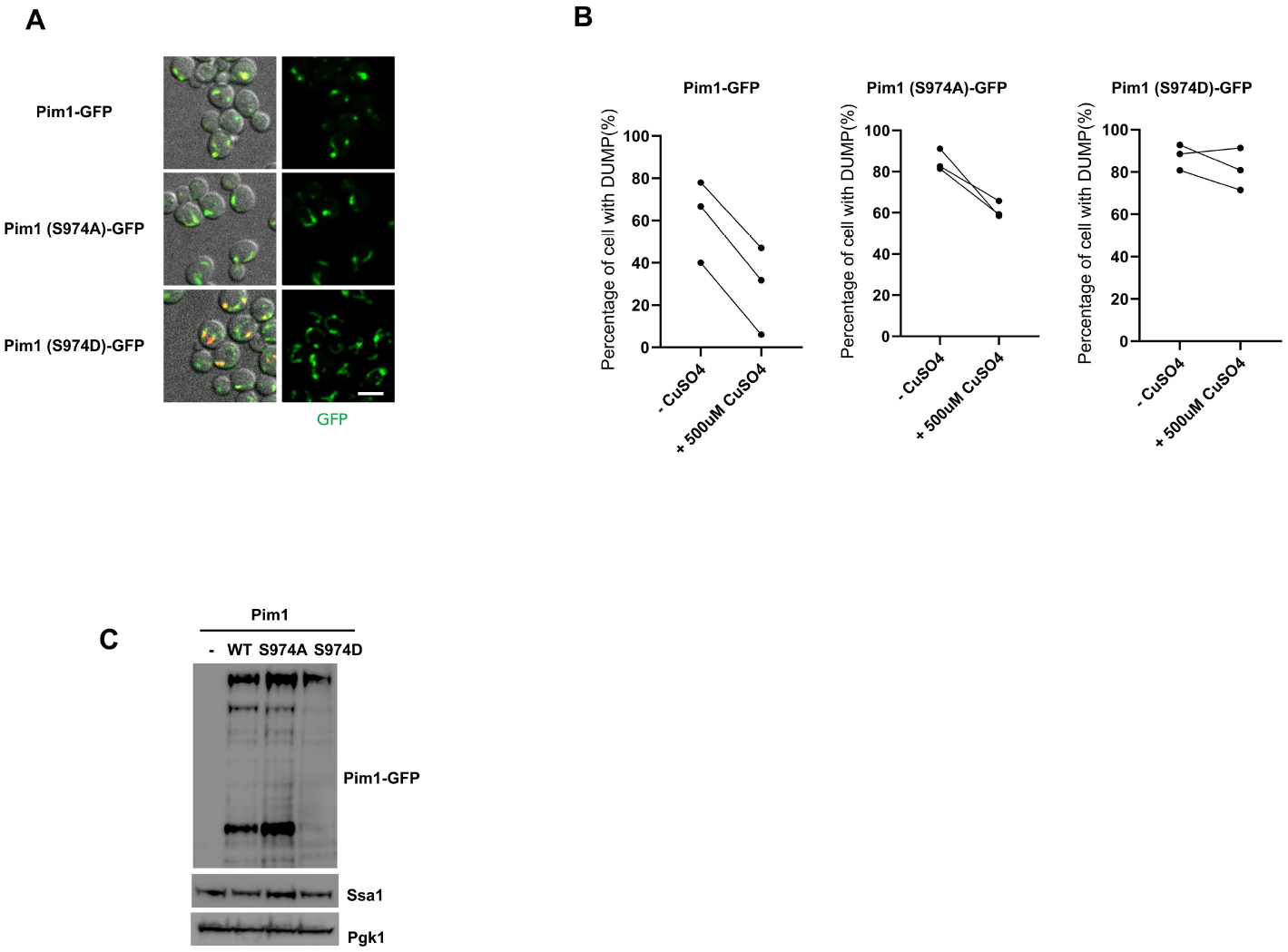
(A). Localization of Pim1 wildtype and the mutants in yeast cells. (B) Quantification of percentage of cell with mitoFluc labeled DUMP structures in (Figure 5G). Paired t-test was used for statistical analysis. (C) Western blot showing the levels of Pim1 wildtype and the mutants in yeast cells.

**Supplementary Figure 3.**
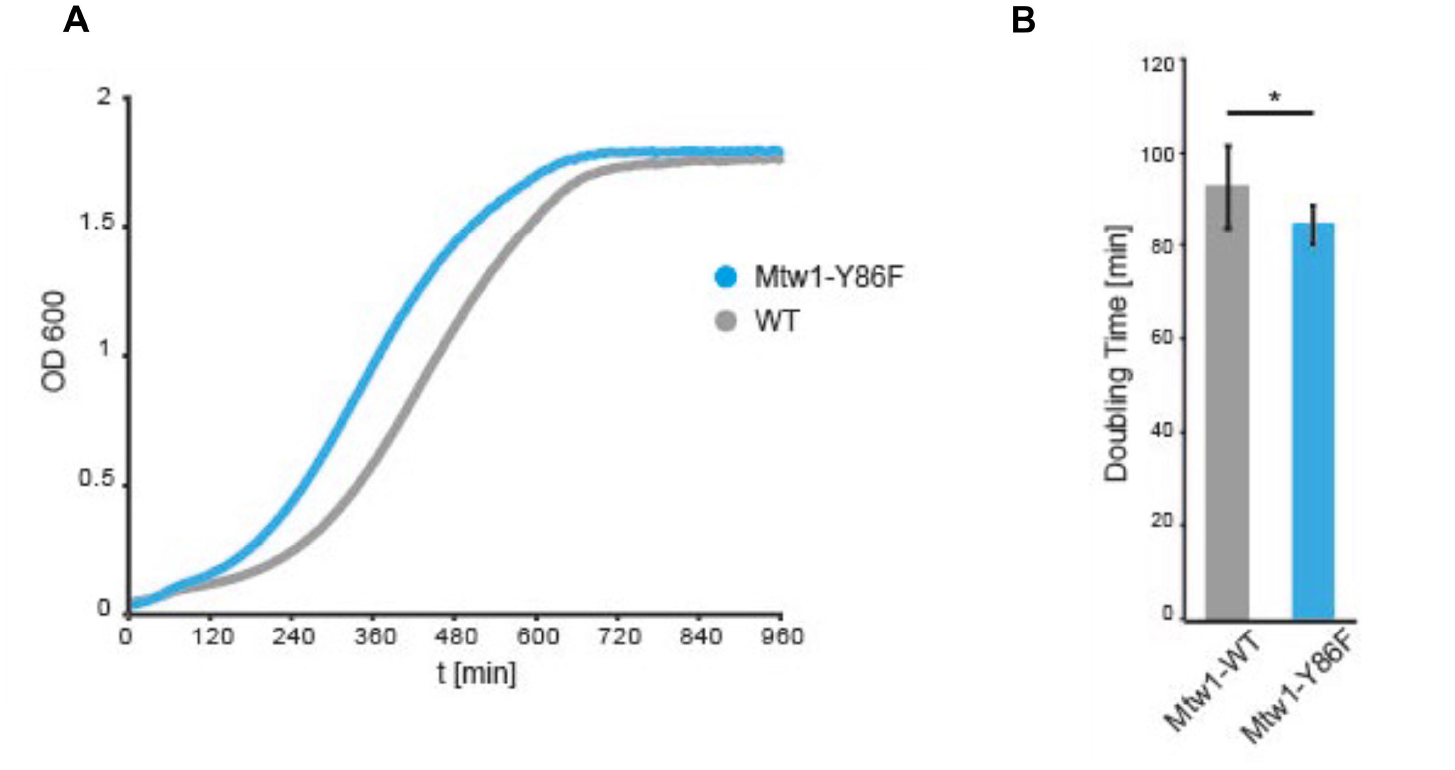
(A) Growth curves of wild-type (BY4741) and Mtw1-Y86F yeast strains. Cells were diluted from an overnight culture to OD_600_=0.03 and the OD_600_ was measured every 5 minutes for 16 hours using a microplate reader. Growth analysis was performed in 10 replicates per strain. (B) Doubling times for wild-type and mutant strains were calculated for each replicate and compared using Student’s t-test (mean=92.97 min, p-value=0.04).

